# The effect of virtual visual scene inclination transitions on gait modulation in healthy young vs older adults - a virtual reality study

**DOI:** 10.1101/2023.03.14.532630

**Authors:** Amit Benady, Sean Zadik, Sharon Gilaie-Dotan, Meir Plotnik

## Abstract

Bipedal locomotion requires body adaptation to maintain stability after encountering a transition to inclined walking. A major adaptation is the change in walking speed to maintain optimal energy consumption. When transitioning to uphill walking, people exert more energy to counteract gravitational forces pulling them backward, while when transitioning to downhill walking people break to avoid uncontrolled acceleration. This behavior is controlled by three main senses: *proprioception* and *vestibular* (aka body-based cues), and *visual* cues. In this study, we aimed to measure the influence of the visual cues on these modulations in healthy older adults and compare it to healthy young adults. To that end, we used a fully immersive virtual-reality system embedded with a self-paced treadmill and projected visual scene that allowed us to manipulate the inclinations of the treadmill and the visual scene in an independent manner. In addition, we measured the visual field dependency of each participant using the rod and frame test. The group of older adults presented the braking (decelerating), and exertion (accelerating) effects, in response to downhill and uphill visual illusions, respectively, in a similar manner as the young group. Furthermore, we found a significant correlation between the intensity of the speed modulation and the visual field dependency for each group separately, however the visual field dependency was significantly higher in the elderly adults. These results suggest that with aging individuals maintain their reliance on the visual system to modulate their gait in accordance with surface inclination in the same manner as young adults.

## Introduction

Nowadays, with the increase in life expectancy, and with older adults becoming larger portion of humanity, there is growing interest in the manifestation of aging in different behavioral domains. Locomotion is one example for a behavioral change that is modified significantly with aging. To control locomotion the motor system utilizes Multi-Sensory Integration (MSI), i.e., processing multiple unisensory inputs into a unique coherent percept. The three main sensory inputs are *proprioception* and *vestibular* cues that are known together as body-based cues, and *visual* cues^1–3^, usually all three are integrated in a synergia to maintain stable locomotion. An important aspect of locomotion is the ability to maintain stability (i.e., balance control) while propelling the body forward. In general terms, a system is deemed to be stable if following a transition, it reverts to its previous or another stable state. The balance control is related to both *stable posture*, which refers to static stability, and *stable locomotion* that refers to dynamic stability. In the older adult population, in comparison to the young population, reduced balance control capacity in both static and dynamic balance control were reported^4^. This reduction is due to several changes that occur with aging and involve alterations in the MSI component weights, specifically increased visual dependency with aging^5,6^. An additional major age-related change is the decrease in gait speed. Gait speed is known to be the most prominent age-related change with spontaneous gait speed decrease of about 1% per year from the age of sixty and afterwards, until reaching less than 1.0 m/s which is considered abnormal^4,7,8^. As walking on inclined surfaces is known to affect walking patterns, and specifically modulate gait speed, it is commonly used to assess stability^9–11^. Two examples for inclined walking adaptations are the exertion and braking effects; during uphill walking, the exertion effect is applied to counteract the gravitational forces pulling backwards, allowing one to maintain a stable walking speed, typically slower than the self-selected speed during leveled walking^9,12,13^. During downhill walking, the braking effect prevents uncontrolled speeding up and allows one to descend in a stable walking speed. In this case there is more heterogenicity among people, as some will increase, and some will decrease their preferred walking speed^9,12,14^. When walking on inclined surfaces, body-based cues are influenced by physical forces exerted on the body, while visual cues are assumed to be influenced by top-down expectations likely based on prior visual experience. Visual field dependence is considered as the level of reliance on visual cues in comparison to body-based cues^15,16^. A popular way to evaluate it is using the rod and frame test^10,17–19^. Visual field dependency varies across healthy people^20,21^, and was found to be higher in individuals with pathologic or physiologic (e.g., aging) balance related conditions^22–26^. To evaluate the contribution of visual cues as part of the MSI, there is a need to artificially manipulate them in an independent fashion from the body-based cues and explore their relative “weight”. To that end, we used a novel paradigm recently presented by our lab using a fully immersive virtual-reality (VR) system where participants walked on a treadmill that was operated in a self-paced mode and were presented with virtual visual scenery projected on a large dome shaped screen. We transitioned the visual scene’s apparent inclination independently of the physical inclination of the treadmill during walking trials and examined the gait modulations following these transitions. Previous studies done in our lab showed that in a young healthy population, while walking on a leveled treadmill, uphill virtually visually simulated transition is followed by a temporary increase in gait speed, while downhill virtually visually simulated transition is followed by a temporary decrease in gait speed^10,11^. Furthermore, the extent of these speed modulations is proportionally related to the degree of virtual visual scene transition in inclinations between −15° to +15°^10^. These visually guided gait speed modulations represent the virtually induced *exertion* and *braking* effects seen in physical uphill and downhill walking, respectively. In addition, we previously found that gait speed modulations were associated with visual field dependency in a young healthy population^10^. In the present study we turned to explore to what extent these effects are preserved in older adults. Since there is growing reliance on vision with aging^27,28^, we hypothesized to see similar locomotor adaptations following virtually visual transitions in an elderly healthy population, but with a higher relative change in walking speed in comparison to the young healthy population. Furthermore, we hypothesized that the visual field dependence would be higher in the elderly population yet associated to their gait speed modulations.

## Materials and Methods

### Participants

Data collected from twelve young healthy participants (mean age ± SD: 26.53±3.09 years old, six males, see Benady et al., 2021^10^) and form fifteen healthy older adults (mean age ± SD: 68.93±4.17 years old, 8 males) were used for this study. Exclusion criteria were physical or visual restrictions (in case needed, they used their daily eyeglasses), cognitive limitations, and any sensorimotor impairments that could potentially affect locomotion or the ability to adhere to instructions. The Institutional Review Board for Ethics in Human Studies at the Sheba Medical Center, Israel, approved the experimental protocol, and all participants signed a written informed consent prior to entering the study.

### Apparatus

The different experimental apparatuses, procedures and outcome measures were elaborately described in our previous work^10,11^. Herein is a brief description:

#### Virtual reality system

Walking experiments were conducted in a fully immersive virtual reality system (CAREN High End, Motek Medical, The Netherlands) containing a moveable platform with six degrees of freedom. A self-paced treadmill was embedded in the moveable platform, allowing participants to adjust the treadmill speed according to their preferred walking speed (Plotnik et al., 2015).

#### VR version of the Rod and Frame Test

The rod-and-frame test was used to assess the visual field dependence for each participant. The test measures how visual perception of the orientation of a rod is influenced by the orientation of a peripheral frame around it. The test was implemented in our lab using Unity software and C# scripting. The participants sat straight in front of a computer screen wearing head mount device (HMD) VR glasses (HTC VIVE, HTC; New Taipei City, Taiwan). They were instructed not to tilt their head or move during the test. The VR environment contained of a central white rod (11° long) with a peripheral frame around it (occupying ~16°*16° of the visual filed) rotated at a trial-specific orientation. The center of the rod and the frame were aligned, but each one with its own independent orientation. Both the rod and the frame were white and presented on a black background (screen resolution was 1920*1080). During the test, a sequence of 28 trials were consecutively presented during which the frame was tilted at one of seven possible random positions: 0/±10/±20/±30 degrees (0 was vertical, + was clockwise), each position was presented four times^20^. For each trial, the rod was at a random orientation (sampled from 0-180 degrees range distribution), regardless of the position of the frame. The participants’ task was to orient the rod upright (i.e., perpendicular) to the true horizon, irrespective of the surrounding frame’s orientation. This was achieved by rotating the rod around its center in either direction using the VR system’s remote control, the surrounding frame was not changed by this manipulation. Once the participants estimated it as being upright, they pressed a button on the remote control, which led to the clearing of the display and the beginning of another trial.

### Procedure

#### VR rod-and-frame test

The rod-and-frame test was the first task that the participants performed in the experiment (after filling the informed consent). Firstly, we made sure that the participant felt comfortable with the head mount device (HMD). Secondly, the participants underwent a short practice trial to confirm that they fully understood the task. Afterwards, 28 test trials were conducted and measured accurately. There was no time limit, and typically the whole session lasted 10 minutes, including the practice trial.

#### Gait trials in large-scale VR system

##### Habituation period to walking in self-paced mode during leveled and inclined surfaces

The participants were attached to a safety harness on the moveable platform during all walking conditions (Figure 1). The first part of the habituation took about 10-15 minutes and was aimed to familiarize the participants with the self-paced mode of the treadmill. During this part, the participants walked at their preferred speed until they felt comfortable, then the experimenter asked them to increase and decrease their speed until they mastered the walking in a self-paced mode. The second part of the habituation included one walking trial of each of the three possible inclinations (i.e., leveled, uphill, and downhill) when the visual and the physical cues were synchronized (‘congruent’ conditions; see more details below). Each trial lasted three to four minutes.

**Figure 1.**
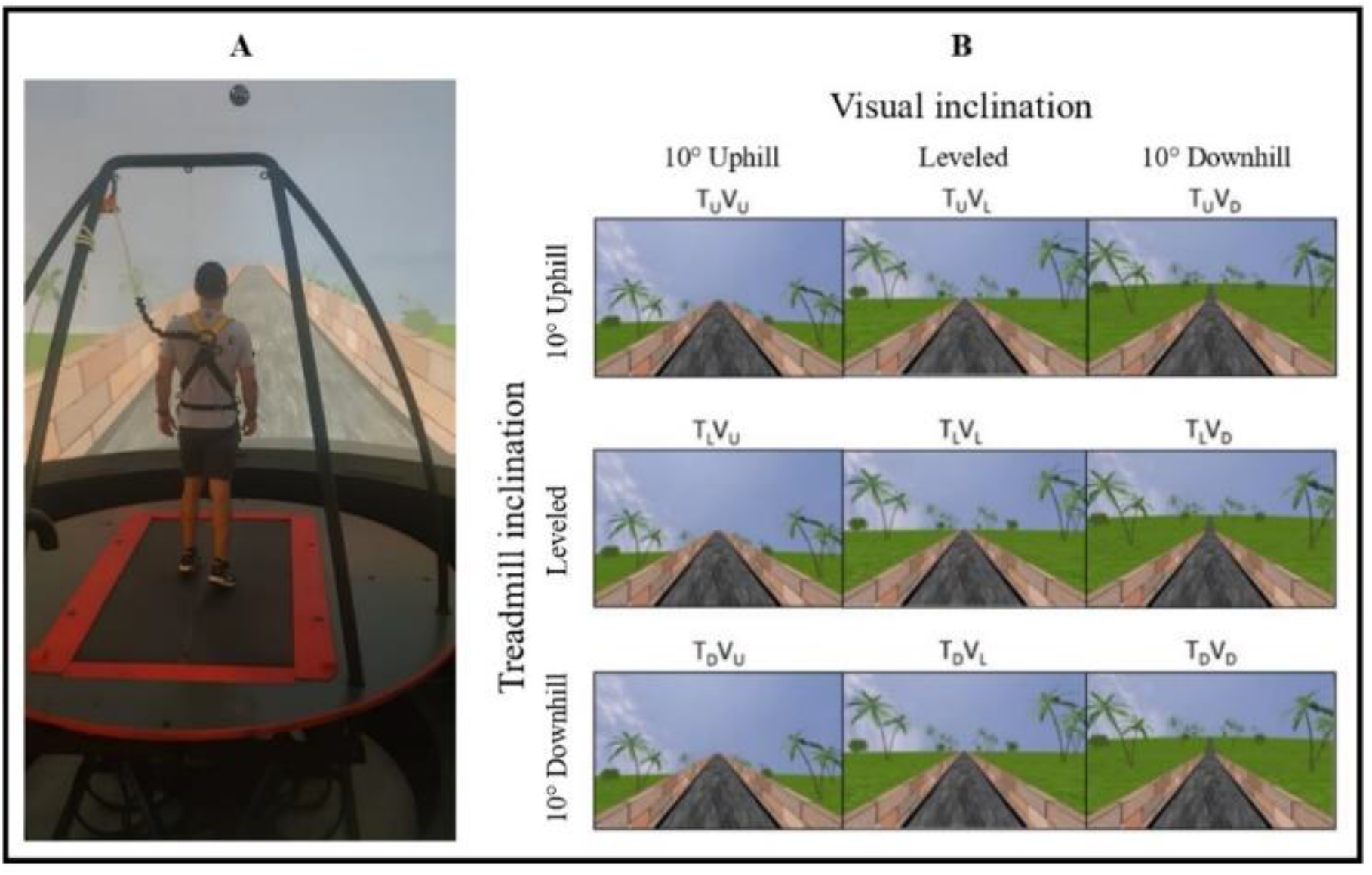
Apparatus (A) and the experimental conditions (B). **(A)** A fully immersive virtual reality system containing an embedded treadmill synchronized with projected visual scenes, wherein this example, the treadmill is leveled, and the vision is uphill (see T_L_V_U_ in B). **(B)** Experimental conditions. Each condition started with leveled walking and only after reaching steady-state velocity (SSV), a transition (5s) occurred to one of nine walking conditions presented in a random order. The conditions include transitions of the treadmill (T) and/or visual scenes (V) to 10° uphill (_U_), remaining leveled (_L_) or to −10° downhill (_D_). Rows depict treadmill inclination changes, and columns depict visual scene inclination changes. Visual scene inclination effect was achieved by the road appearing above (uphill), below (downhill), or converging (leveled) with the line of the horizon. In addition, the peripheral greenery is exposed more (downhill) or less (uphill) by the road. Vision-treadmill congruent conditions appear on the diagonal (T_L_V_L_ for continued leveled walking, T_U_V_U_ and T_D_V_D_ for uphill and downhill walking, respectively). Vision-treadmill incongruent conditions below the diagonal represent conditions with visual scene inclination more positive than the treadmill’s (T_L_V_U_, T_D_V_U_, T_D_V_L_), and above the diagonal visual scene inclination more negative than the treadmill’s (T_U_V_L_, T_U_V_D_, T_L_V_D_). See Methods for more details. (Adopted from Benady et al^10^.

##### Gait experiments

The participants were informed that they would perform several short gait trials with short intervals between them. They were instructed to walk “as naturally as possible” and that “inclinations may occur during walking”. Each walking condition started with the participant in a standstill position and then progressed into walking with both the treadmill and the visual scene leveled until reaching steady-state velocity (SSV) for 12 seconds. Once reaching SSV, a 5-second-long transition of the treadmill and/or visual scene occurred (except in the congruent leveled condition, where no actual transition occurred). Post transition, each condition lasted another 65 seconds, until the treadmill slowed down and stopped altogether.

##### Experimental conditions

The protocol included nine walking conditions that the participant encountered in random order. Inclination of the treadmill (T) and/or visual scenes (V) transitioned to 10° uphill (U), remained leveled at 0° (L) or transitioned to −10° downhill (D). Figure 1 shows the 3×3 experimental design, where rows represent treadmill (T) inclination and columns represent visual scene (V) inclination. Congruent conditions where the treadmill and visual scene inclinations are synchronized were set as baselines. These conditions are present on the diagonal of Figure 1; upper left-uphill (T_U_V_U_), middle-leveled (T_L_V_L_), lower right-downhill (T_D_V_D_) walking. Treadmill-vision incongruent conditions include the following visual scene manipulations: for treadmill uphill inclination, the vision was leveled (T_U_V_L_) or downhill (T_U_V_D_), for treadmill leveled inclination, the vision was uphill (T_L_V_U_) or downhill (T_L_V_D_), and lastly, for treadmill downhill inclination, the vision was leveled (T_D_V_L_) or uphill (T_D_V_U_).

### Outcome measures

#### Gait speed related variables

To assess the post-transition effects on gait speed, we looked at (i) the magnitude of the peak/trough of gait speed relative to the SSV (%); and (ii) the time of this peak/trough after the transition (seconds). We refer to the transition start time as time zero (t=0).

##### Steady-state velocity

A real-time algorithm monitoring treadmill speed determined the SSV. According to the algorithm, SSV is attained after (i) minimum 30s of walking from the beginning of the trial, and (ii) a consecutive period of 12s with gait speed coefficient of variance less than 2%. Upon satisfying both conditions, the transition of the treadmill and/or visual scene inclination (as appropriate for the experimental condition) was automatically triggered.

##### Normalization of gait speed

Normalization of gait speed for each participant in each experimental condition consisted of three steps: (i) gait speed was divided by the averaged SSV at every second, (ii) the ratio between gait speed and SSV was presented as a percentage, and (iii) the normalized trace was shifted so that the mean value of the SSV period would be zero. Following these steps, it was clear to distinguish between the responses of increased and decreased speed following the transition.

#### Standardized response to virtual inclination

To compute this index, we used data from the incongruent T_L_V_U_, T_L_V_D_, T_D_V_U_, T_U_V_D_ walking conditions. The averaged absolute values from these four conditions of the peaks/troughs (%) relative to the SSV were calculated for each participant.

#### Visual field dependence index

In each trial of the rod and frame test, the degree of deviation of the rod from the true upright position was recorded as the position error. For each participant, the mean position error of the seven different frame angles was calculated. Data from all participants were grouped by the frame angle^20^. We defined the visual field dependence index as the average position error when the frame was projected at ±20 degrees (8 trials in total, 4 trials of +20° and 4 trials of −20°). This parameter allowed us to evaluate individual differences in visual field dependence.

### Calculation of Ratio of Gravity-Induced Behavior

To assess the amount of influence of gravity on walking speed (WS), we calculated the Area Under the Curve (AUC) second-by-second (i.e., at 1 s, at 2 s… at 60 s) from V(t) (i.e., 9.8*sinθ*t) and WS for congruent uphill and downhill conditions. We defined the ratio according to the following equation (we added a multiplier of 100 to avoid extremely small numbers):

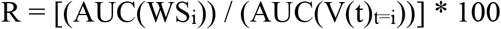

The index “i” refers to the time (in seconds) post-transition. The ratio quantifies the degree to which WS approximates the velocity of a free body. A positive ratio indicates that both WS and a free body accelerate/decelerate at the same direction (i.e., both accelerate when downhill and decelerate when uphill). A ratio further from zero suggests more gravitational influence on walking.

### Linear summation model

Given that locomotion is maintained by the IMG, we used a linear weighted summation model to estimate the weight of visual and body-based cues^1,11^. The general model is presented in the following equation:

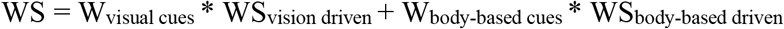

WS is defined as the integrated behavior after considering the weight of each component. W is defined as the weight of a unimodal cue (*visual* or *body-based*). For simplicity purposes, herein we will explain the linear summation model for uphill walking, the same concept but in the opposite direction accounts for the downhill walking. We assume that that WS in condition TLVU (WS_vision-up_) is driven exclusively by uphill visual cues. Contrariwise, we assume that WS in condition T_U_V_L_ (WS_body-up_) is driven exclusively by the body-based cues. The weight of each unimodal cue can be calculated using the following equations. As expected while assuming that vision and body-based cues are the main factors influencing on locomotion, it is seen that W_vision_ + W_body_ =1.

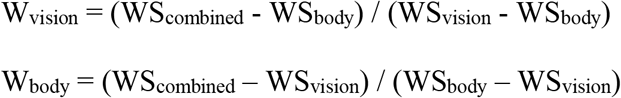

Finally, WS_up_ represents the congruent condition T_U_V_U_ and can be calculated using the following equation:

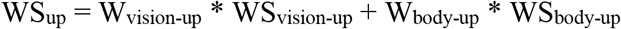

### Statistical analyses

Values are presented by their group mean values (± SE). Differences in the magnitude and timing of normalized walking speed in the incongruent walking conditions (i.e., T_L_V_U_ and T_L_V_D_) between the young and older adult groups were compared at the peak/trough of change, using a two tailed t-test, with a p-value of equal or less than 0.05 indicating a significant difference. Similarly, the difference between the rod and frame index of both groups was compared with the same method. The after effect, comparing the return from the peak/trough to the SSV was calculated by a One-way t-test with 0 (i.e., SSV) as the index. Pearson correlation coefficient was computed to evaluate the relationship between visual field dependence index and (i) the standardized response to virtual inclination and (ii) the age of participants. All data were analyzed using SPSS software (v25, IBM).

## Results

### Gait speed modulations following physical and/or virtual inclination transitions

We initially compared the behavioral change measured by gait speed modulations in the young vs older adult’s groups following a virtual (i.e., visual) and/or physical (i.e., treadmill) inclination transitions. Figure 2 depicts the averaged normalized walking speed relative to the SSV for each condition, one minute pre- and post-transition. For the treadmill up conditions there was a gradual monotonic change in speed in both groups, with a more prominent effect in the older adult group (see Figure 2, first row how the purple (older adult group) line is lower (i.e., slower) than the orange (young group) line). For the treadmill leveled conditions, the same patterns were seen between the groups for all three visual scene inclinations. When the visual scene remained leveled (T_L_V_L_), both groups maintained roughly the same speed as the SSV. When the visual scene transitioned upwards (T_L_V_U_-i.e., virtually induced exertion effect), both groups temporarily increased their walking speed until reaching a peak. No significant change was seen neither in the time of the peak (mean ±SE: young=10.37±0.64sec, older adults=11.05±1.34sec, p=0.65), nor in the amplitude of the peak (young=18±3%, older adults =12±4%, p=0.25). Note that after the peak, the older adult group returned to their SSV, while the young group decreased their speed until a new steady state, which was higher than their previous SSV. Regarding the opposite condition, where the treadmill remained leveled, but the visual scene transitioned downward (T_L_V_D_-i.e., virtually induced braking effect), both groups decreased their speed until reaching a trough with no significant difference in the peak’s time (young= 8.08±0.59sec, older adults =8.06±0.48sec, p= 0.98), nor between the magnitude of both groups (young=25±4%, older adults =32±6%, p=0.32). After the trough, the young group returned to their previous SSV after 14.4±0.32sec while the older adult group returned after 19.68±0.54sec (p=0.02). Interestingly, in the treadmill down conditions both groups showed opposite walking patterns; in the congruent condition TDVD, while the young group constantly increased their walking speed, the older adult group initially braked and decreased their speed, with a monotonic increase during the next minute after the transition. This pattern was roughly the same also in the T_D_V_U_ and the T_D_V_L_ conditions.

**Figure 2.**
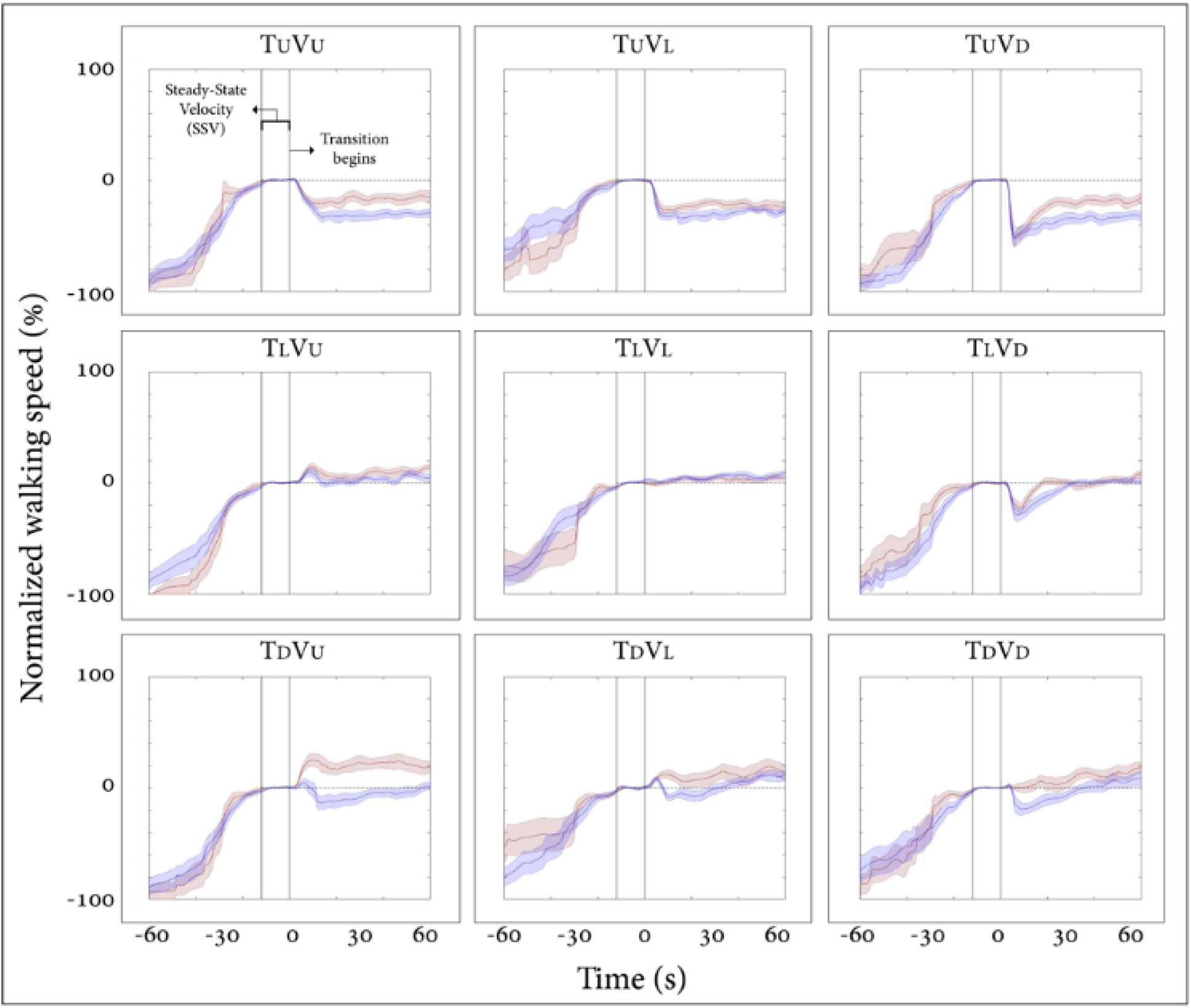
Normalized walking speed in the young (N=12) and older adults (N=15) groups. X-axis represents the time (s); 60 seconds before and after the transition, y-axis represents the average normalized self-paced gait speed relative to steady-state velocity (%) for each condition. Orange lines represent the young group and purple lines the older adult group, shaded colors represent the standard error for each group. Time zero demarcates the end of the steady-state velocity period, after which a 5s transition of the treadmill and/or visual scene occurred. Conditions: treadmill (T) and/or visual scenes (V) inclination transitioned to 10° uphill (U), remained leveled at 0° (L), or transitioned to −10° downhill (D). Note that there was no difference in speed modulation patterns or magnitude between both groups following the visually induced *braking* and *exertion* effects.

### Relation between visual modulation of gait speed during visual-physical incongruent conditions and visual field dependence

In order to further test our findings from our previous study^10^ that showed a significant correlation between the magnitude of visual modulation on gait speed during virtual inclination changes and individual’s visual field dependency in a young group, we now computed these measures for the older adult group. We calculated for each participant (i) the magnitude of visual modulation on gait speed based on the gait speed changes in the incongruent conditions (i.e., standardized response to virtual inclination, see Methods for calculation), and (ii) the index of visual field dependence estimated by the rod and frame test (see Methods). As expected, we found also in the older adult group a significant correlation between the standardized response to virtual inclination and the index of visual field dependence estimated by the rod and frame test (c.f. Methods), (Fig. 3, upper panel; blue circles, Pearson’s r=0.76, p= 0.001). For comparison the data from young healthy adults were plotted on the same graph (orange points) also reflecting significant correlation between these two measures (Fig. 3, upper panel; orange circles, Pearson’s r=0.76, p= 0.004). No correlation difference was seen between both correlations (p=0.5). These findings suggest that both parameters rely upon associated mechanisms across lifespan. Furthermore, a significant correlation was seen between the rod and frame index and the age of the participant only in the older adult group (Figure 3, lower panel; Pearson’s r(13)=0.67, p=0.006), but not in the young group. In addition, a t-test showed significant difference (p<0.001) between the rod and frame indices of the two groups (young vs older adults).

**Figure 3.**
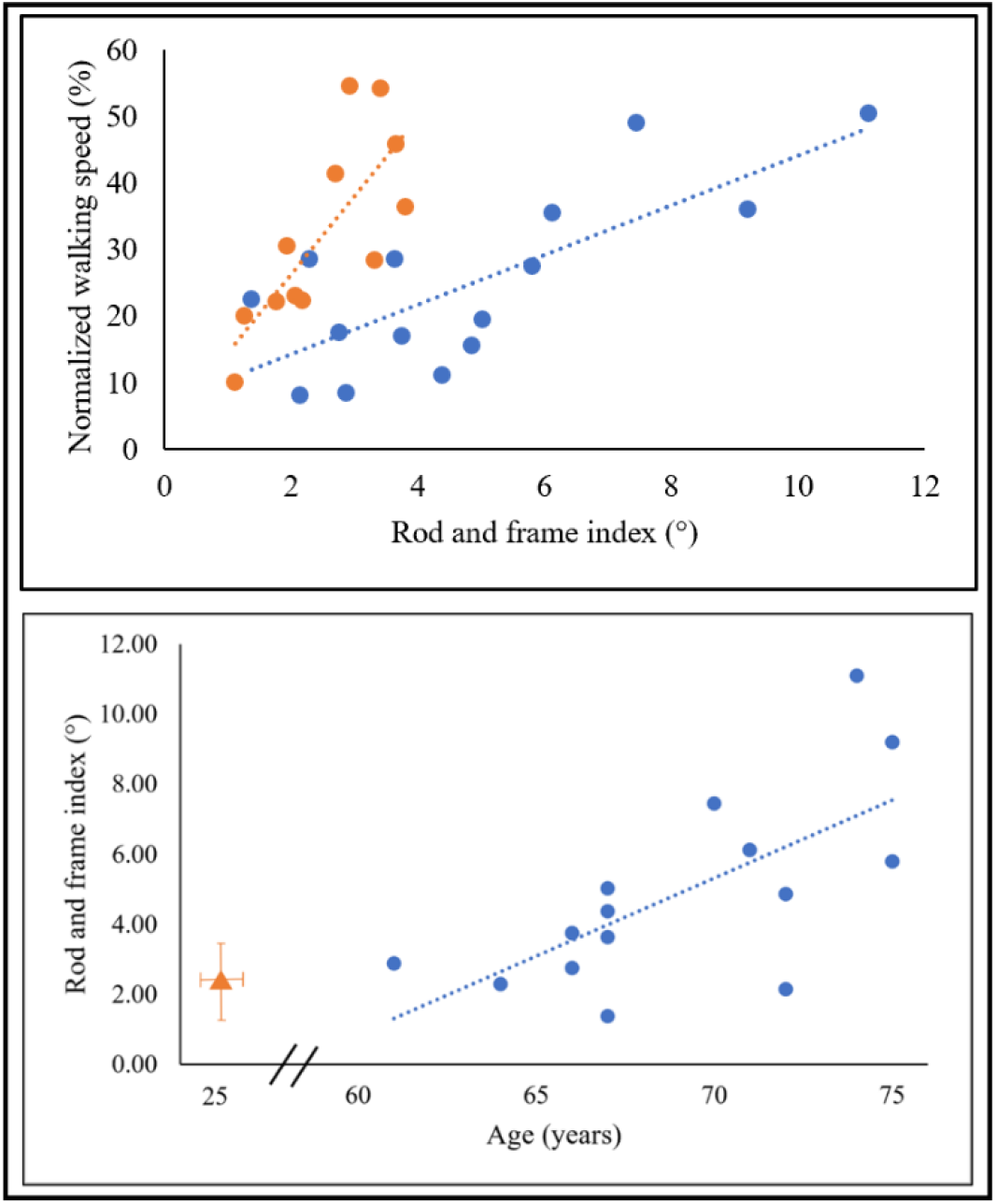
Upper panel. Averaged standardized response to virtual inclination relative to the visual field dependence index. Blue circles represent the older adult group (N=15) and orange circles represent the young group (N=12), each circle represents one participant. X-axis represents the visual field dependence as assessed by the psychophysical rod and frame test while seated. Y-axis represents the standardized response to virtual inclinations based on the treadmill-visual incongruent walking conditions. A significant correlation was seen between these two values for each group independently (young: Pearson’s r=0.76, p= 0.004), (older adults: Pearson’s r=0.76, p= 0.001). **Lower panel. Visual field dependence index relative to participant’s age.** The extent of visual field dependency during locomotion in incongruent conditions is correlated with the age of the participant in the older adult group, but not in the young group. X-axis represents the age of the participant (years), y-axis represents the rod and frame index. Blue circles represent the older adult group (N=15) and orange circles represent the average of the young group (N=12) for reference. A significant correlation was found in the older adult group (r(13)=0.67, p=0.006).

### Ratio of gravity-induced behavior in congruent uphill and downhill walking

To further compare between the two groups, we measured the braking end exertion effects post-transition over time. For this, we computed the normalized ratio between the areas under the curve of gait speed for the congruent uphill and downhill walking conditions, divided by free body velocity at the same inclination (i.e., V(t)=g*sin±10*t; where g is the gravitational constant) (Figure 4). The higher the ratio (i.e., further from zero), the stronger the effect of gravity and the weaker the exertion and braking effects are. The analysis revealed a differential response to gravity forces between both groups, both in the uphill and downhill walking (see statistical results in figure legend). For uphill walking, the same pattern was seen between both groups (c.f. Figure 2, upper left corner), but with a greater ratio in the older adult group, reflecting a greater influence of gravity with a smaller exertion effect. The turning point, which demarcates the time (in seconds after the transition), when the exertion effect begins, and participants start to counteract gravity forces, was similar in both groups (T_Uphill_=10s). For downhill walking, the older adults group showed initially the same pattern of a monotonic decrease in walking speed, with eventually a gradual return to SSV. Their turning point was almost the same as in the uphill condition (T_Downhill_=11s), meaning that in both inclinations, after 10s-11s the older adults group applied the exertion effect to counteract gravity. The walking pattern for the young group was initially maintaining SSV with a gradual increase in speed. Their turning point was at 9s, in this case reflecting a decrease in the braking effect which prevents an uncontrolled acceleration forward, followed by a gradual monotonic increase of the walking speed (c.f. Figure 2, right lower corner).

**Figure 4.**
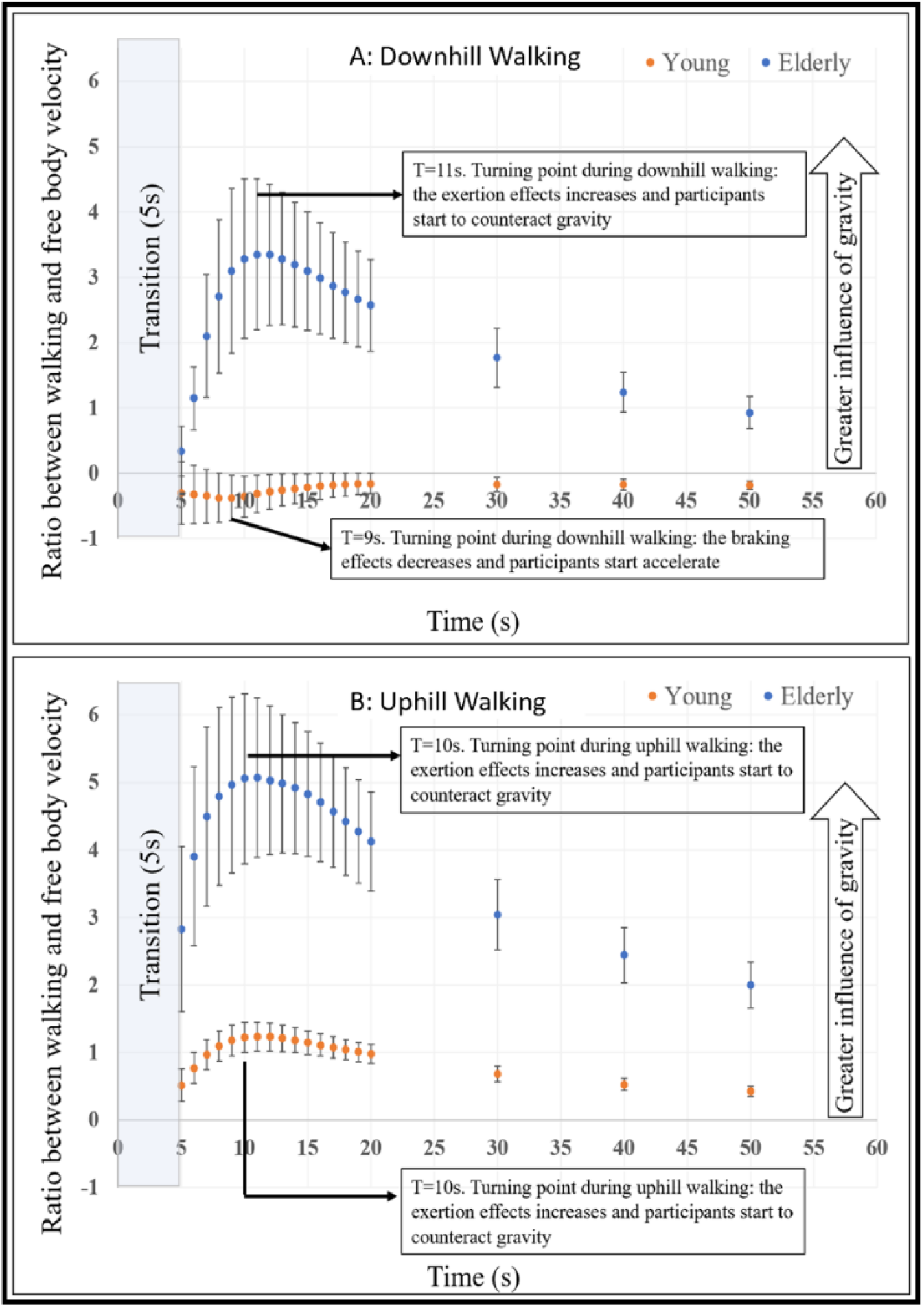
Ratio of gravity-induced behavior in congruent uphill and downhill walking. The average ratio between participants gait speed (WS) and velocity of a free-moving body (V(t)) over time for ±10° downhill (upper panel) and uphill (lower panel) walking. Error bars represent the SE. The turning points represent the time (seconds) in which the participants either applied or rejected the exertion or braking effects, to counteract or engage gravitational forces during walking. A statistical difference (p<0.05, corrected for multiple comparisons K=19) was seen between the young and older adults only after the turning points.

### Application of the linear summation model for congruent uphill walking in the young vs older adult groups

We applied the linear summation model (see Methods) to evaluate the weight of the *visual* cues in comparison to the *body-based* cues over time during uphill walking (Figure 5, upper panel), and to further examine if our prediction indeed fits the true behavior during congruent uphill walking (Figure 5, lower panel). This model assumes that true uphill walking is the integration of the unimodals contributing to it, in our case visual (vision-up) cues and body-based (treadmill-up) cues. Each cue has a relative contribution for every time-period, but at any time the sum of all unimodals equals one. This model enables estimation of the sensory reweighting of the IMG components. For example, a weight of body-based cues near 0 indicates that at the same time-point, locomotion predominantly relies on vision. Our model shows the relative weight of each unimodal cue as we continue walking. This model did not fit our participants for downhill walking as it is known that in young people walking modulations are less homogeny during downhill walking as there are people who increase and people who decrease their speed^9,12,14^, while older adults usually decrease their speed^29^. In contrast, uphill walking has a more homogeny pattern with most people decreasing their speed, both in the young and older adult population^29,30^.

**Figure 5.**
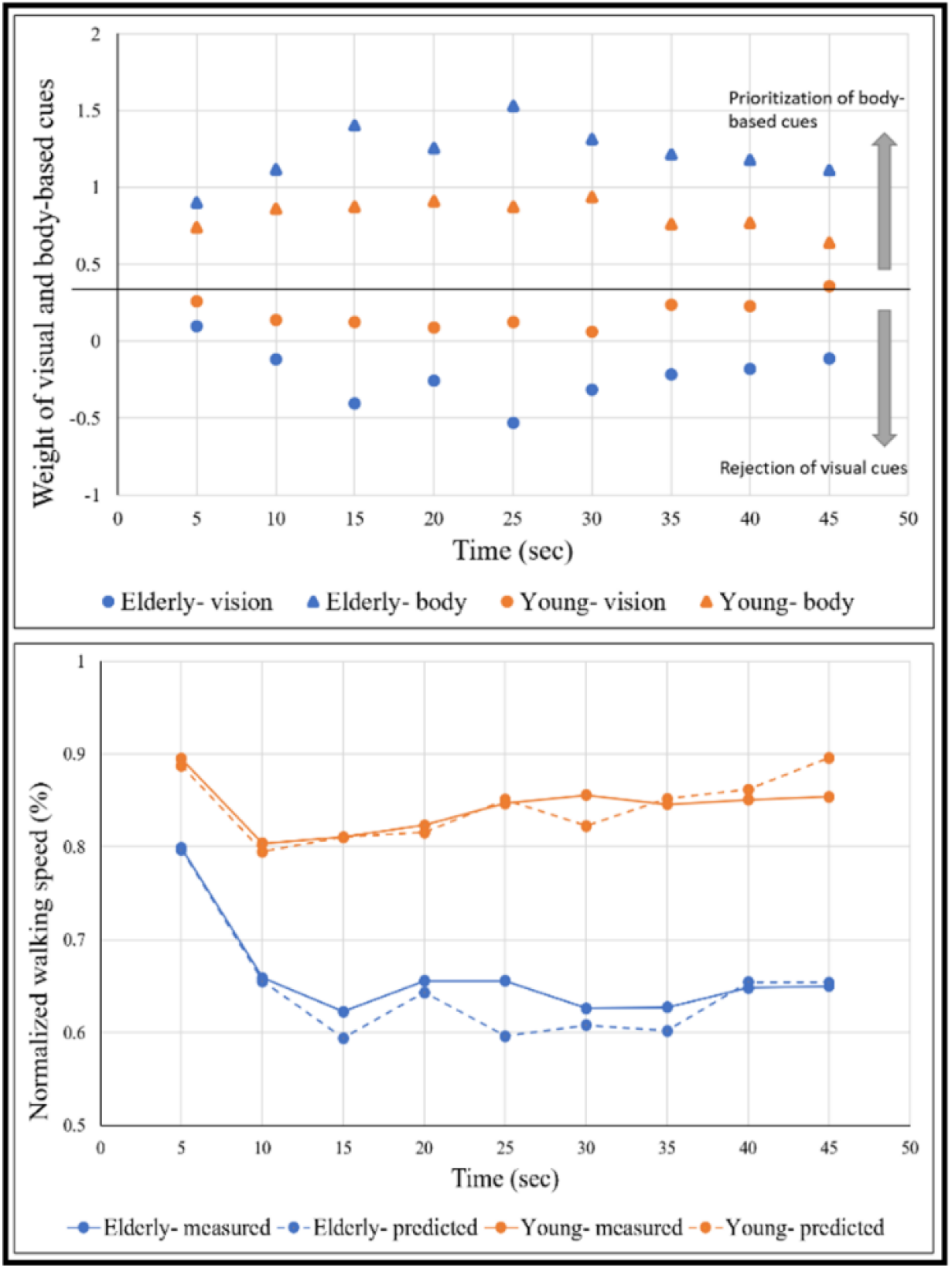
Congruent uphill walking. (Upper panel) - Sensory reweighting. (Lower panel) - Linear summation model (see Methods). X-axis depicts the time (sec), time zero demarcates the beginning of the transition (5s long). Blue color represents the older adult’s group and orange color represents the young group. **Upper panel.** The weight of visual and body-based cues (proprioception and vestibular) during locomotion over time. Circles represent the weight of visual cues, triangles represent the weight of body-based cues. **Lower panel.** Comparing our measured gait speed to the expected gait speed according to the linear summation model over time in congruent uphill walking (c.f. Figure 2, upper left corner). Solid lines represent the measured walking speed and dashed lines represent the expected gait speed.

## Discussion

In this study, we compared the weight of visual cues to locomotion modulation under physical and/or visual changes between healthy young and older adults. We manipulated vision independently of body-based (proprioception and vestibular) cues and measured the changes in walking speed. We found that the same pattern of walking adaptation to virtually induced braking and exertion effects (either slowing down or speeding up, respectively) is employed by both groups. Group measures in the magnitude and in the timing of the peaks/ troughs of the braking and exertion effects were not significantly different. However, we did see a change in walking speed modulation in incongruent condition when the treadmill was transitioned downward. We also observed a significant correlation between the magnitude of the virtually induced braking and exertion effects, and the individual’s visual field dependency as measured by the rod and frame test for each group separately, but with significantly bigger reliance on visual field in the older adult group. Finally, we applied the linear summation model during uphill walking, which showed the relative weight of each unimodal cue as we continue walking in both the young and older adult groups and found the expected speed pattern fit the real walking speed pattern as measured in the congruent uphill condition.

### The relative weight of visual cues during physical and/or visual manipulations

#### Treadmill leveled conditions

When walking with the treadmill leveled, both groups showed the same pattern of gait speed modulation for all three virtual visual inclinations (c.f. figure 2, middle row). As older adults have increased reliance on visual cues^27,28^, an observation that was confirmed in our study as can be seen by the results of the rod and frame test (Figure 3) we expected to see larger speed modulations following virtually induced braking and exertion effect. Surprisingly, no significant difference was seen in the magnitude nor the timing of the peaks/troughs. Lack in differences might be related to the cohort of elderly adults who participated in the study. While being above the age of 65, most of the participants were relatively fit, possibly reflected by the fact that they consented to participate in the study after being informed that a 10-degree inclined walking may be required. Thus, our study might be limited by the fact that the older adults of our study might not be representative for a typical ~70-year-old individual. We did not encounter young adults declining to walk at 10 degrees. This finding strengthens the notion that MSI deterioration is not affected primarily by aging, but rather by physical condition and lifestyle, factors that can be controlled and modified. At the same time, our study convincingly observed that while maintaining the behavioral constructs of braking and exertion effects, older adults resist the consequences of the gravitational forces upon their walking speed to a lesser extent than young adults do. This fact was reflected by the ratio of gravity-induced behavior. We believe that this finding is reflecting the general reduction in volition of locomotion that is also expressed by additional age-related modulations of speed walking^4,7,8^.

#### Treadmill uphill/downhill conditions

When the treadmill transitioned upwards (c.f. figure 2, upper row), both groups have showed a steady decrease in walking speed across the whole trial, and as expected the older adults walked slower than the young group. Interestingly, when the treadmill transitioned downward both groups showed different walking patterns (c.f. figure 2, lower row). In the congruent T_D_V_D_ condition, while the young group increased their gait speed, the older adults initially applied the braking effect, possibly until they felt in control, and only then let the gravity force accelerate their speed. The deterioration of the multi-sensory integration with aging is seen nicely when the treadmill transitioned downward, and the visual scene remained either leveled or transitioned upward. It appears that an initial increase in gait speed is followed by a decrease in gait speed, and only after about 20 seconds, a steady gait speed is reached. We suggest that the phenomenon is seen primarily when walking downwards, because of an a priori perception that walking upward is less ‘frightening’ and the physical restrictions seen with aging govern on the discrepancy in perception, which eventually is causing a decrease in walking speed. In contrast, while walking down one fears to lose control and fall so the multi-sensory integration is more tuned to maintain stability.

#### Inter-participant variability

In the healthy population, there is a well-established relationship between subjective visual vertical (SVV), which is thought to indicate visual field dependency^17,19^, and postural stability^22,31^. However, the locomotive reactions, which are behaviorally reflected by variations in gait speed, are not fully understood. Herein we strengthen the notion that visual field dependency increases with aging^27,28^, this can be seen by (1) significant increase in the older adults group compared to the young group, and (2) only within the older adults, there is a significant correlation between age and rod and frame index. Future studies should measure visual field dependency across the lifespan to detect at which age the increase starts, and whether is plateaus eventually. The correlation between age and rod and frame index strengthens our previous findings^10,32^ that visual dependence as measured independently by two orthogonal tests, i.e., subjective spatial computerized test, and behavior-based gait speed adaptations reflect the fact that assessment of gravity direction (rod and frame) and consequences (speed modulations) are interrelated in the relevant neuronal pathways. We posit that this finding may hold translational significance, for example, as a new evaluation approach that combines a short walking trial in a visual conflict paradigm with the rod and frame test can potentially estimate visual dependency in locomotion. This may aid in the identification of those who may benefit from visual conflict paradigms, e.g., for gait rehabilitation purposes allowing for a more individualized rehabilitation approach.

#### Limitations

There are several limitations to our study. First, our study included a relatively small sample size, this is more profound within the older adults, as the variation across individuals is higher. Second, using a VR platform may not reflect real life behavior, even more in the older adult group that are less connected to novel technologies and may need a longer habituation period to feel comfortable with the paradigm. Third, we aimed to evaluate the weight of visual cues compared to body-based cues while walking on inclined surfaces. Therefore, we used a self-paced treadmill with a fully immersive projected visual scene, which was shown to have high ecological validity^33^. However, transitions to inclined walking were presented swiftly and within a time window of 5 seconds, while in real life, the individual can anticipate the inclination in advance, based on visual inputs from the environment. This could affect more the older adults, which have decreased vision acuity, and they might encounter the transition at a later stage^34^. Despite this limitation, the results of the present paradigm clearly discern between the roles of vision and body-based cues in adjusting gait behavior with reference to inclinations.

#### Future directions

Overall, uni-senses and body functions are known to deteriorate with aging, such as vision^34^, joint mobility^35^, muscle force^36^ and balance^37^. However, age-related changes of multi-sensory integration is less studied. The changes in performance in MSI tasks may be more sensitive and aid as an earlier predictor to detect future deterioration in daily living activities compared to assessing each uni-sense by itself. Future studies should investigate among other aspects, also the effects of aging on MSI in relation to the autonomic nervous system (ANS) (e.g., heart rate variability). Normal aging is accompanied by a series of complex alterations in the autonomous control of the cardiovascular system, with an increase in the cardiac sympathetic nervous system tone and withdrawal in the parasympathetic nervous system. Although paradigms of incongruent feedback information provide some insight about the mediating effects of external performance cues on perceptual experience, very little is known about the mechanism that controls such behavior, and how it is manifested in the activity of the ANS, especially in older adults. Better understanding these mechanisms can further improve the early detection and clinical treatment outcomes of aging.

## Acknowledgements

The authors wish to thank Ms. Adi Lusting and Mr. Yotam Hagur-Bahat for technical support.

## Author contributions

AB was the main author of this manuscript. AB and SZ recruited and conducted the experiments. AB analyzed the data. SGD and MP supervised the experiments and data analyses as well as critically reviewing the manuscript. All authors approved the manuscript before submission.

